# EREFinder: Genome-wide detection of estrogen response elements

**DOI:** 10.1101/282533

**Authors:** Andrew P. Anderson, Adam G. Jones

## Abstract

**Motivation:** Estrogen response elements (EREs) are specific DNA sequences to which ligand-bound estrogen receptors (ERs) physically bind, allowing them to act as transcription factors for target genes. Locating EREs and ER responsive regions is therefore a potentially important component of the study of estrogen-regulated pathways.

**Results:** We tested and demonstrated the ability of EREFinder, a novel algorithm we developed, to locate regions of ER-binding across the human genome and show that these regions designated by the program occur more frequently near estrogen responsive genes. EREFinder can handle large input files, has settings to allow for broad and narrow searches, and provides the full output to allow for greater data manipulation. These features facilitate a wide range of hypothesis testing for researchers and make EREFinder an excellent tool to aid in estrogen-related research.

**Availability and Implementation:** Source code and binaries freely available for download at https://github.com/JonesLabIdaho/EREfinder, implemented in C++ and supported on Linux and MS Windows.

**Supplemental Materials:** R scripts can be found at https://github.com/JonesLabIdaho/EREfinder

## Introduction

Estrogen receptors (ERs) are the most primitive steroid receptor in the chordate lineage (Thornton et al. 2003) and play a part in a wide range of biological processes within vertebrates. While many studies on ERs have focused primarily on pathologies and medical applications (Arnal et al. 2017, Jia et al. 2015), the pleiotropic effects of ER activity can be wide-ranging (McDonnel and Norris 2002). Consequently, the genome-level details of estrogen signaling, especially with respect to ER binding and its effects of transcription, are topics of considerable interest to a wide range of disciplines, from human health to ecology and evolution. Many factors contribute to the control of gene regulation by ERs, including the type of ligand (Anstead et al. 1997, Arnal et al. 2017), the presence of appropriate cofactors (Arnal et al. 2017, Glass and Rosenfeld 2000), and the genome-level distribution of estrogen response elements (EREs).

The consensus ERE is a palindromic sequence of two half-sites with a 3-base-pair spacer (AGGTCAnnnTGACCT) that is preferentially bound by the ER (Klein-Hitpass et al. 1988, Boyer et al. 2000) making EREs a critical point of contact between ERs and the transcription of ER-regulated genes. Point mutations deviating from the consensus negatively affect the strength of ER binding (Tyulmenkov and Klinge 2001, Geserick et al. 2005, Deegan et al. 2011), and weaker ER binding has been shown to negatively affect transcription of target genes (Klinge et al. 2000, Tyulmenkov and Klinge 2001). Of particular importance are the half-sites (AGGTCA), because strong binding of an ER requires at least one perfect half-site (Tyulmenkov and Klinge 2001, Deegan et al. 2011). Moreover, the number of available EREs in a given region positively correlates with expression of the target gene, whether those EREs are perfect or imperfect (Martinez and Wahli 1989, Kato et al. 1992, Geserick et al. 2005). While some genes are regulated by distant EREs, the larger share of gene regulation occurs via EREs that lie in close proximity to the transcription start site (Carroll et al. 2006, Lin et al. 2007).

Identifying functional EREs has been accomplished by finding estrogen responsive genes and transfecting upstream regions to determine sequence responsiveness (Klein-Hitpass et al. 1988, Martinez and Wahli 1989, Kato et al. 1992) or by employing ChIP-seq protocols (Carroll et al. 2006, Lin et al. 2007). One limitation of these methods is that EREs can only be detected if they are bound by ERs in the given tissue at the given ontogenetic stage, while the remainder of potentially ER responsive genes in the genome could be overlooked. Additionally, EREs that are not bound have the potential to become bound given the correct cofactors and thus might be evolutionarily important but undetected in empirical studies of ER binding. Hence, an *a priori* bioinformatic method to identify potential regions of ER binding in a genome sequence would be a useful complement to current empirical approaches. A handful of software packages already perform this function to some degree either by identifying transcription factor binding sites generally (Frith et al. 2003, Kel et al. 2003.) or EREs specifically (Bajic et al. 2003). However, these programs have shortcomings that render them difficult to apply on a whole-genome basis in the search for EREs, such as small file size imputs or abbreviated outputs that do not display all possible information gathered from a search.

Here we present EREfinder, a novel program for locating putative EREs. EREFinder utilizes the equations developed and empirically validated by Tyulmenkov and Klinge (2001), which predict the binding affinity (*K*_*d*_) of ERα and ERβ to a given DNA sequence. By utilizing a sliding window defined by the user, EREFinder can function both as a detector of EREs and as an indicator of the average expected ER-binding affinity in a given subset of the sequence. We also compare the functionality of EREFinder to two other commonly used ERE detection programs (Frith et al. 2003, Kel et al. 2003.). Finally, we demonstrate EREFinder’s ability to find significant regions in the human genome adjacent to genes that are known to be estrogen responsive.

## EREFinder

Following the formulae presented by Tyulmenkov and Klinge (2001), EREFinder performs a scan across the entire fasta-format input file, evaluating binding affinities of either estrogen receptor α or β. Because the canonical ERE is palindromic, the program reads in one direction along a given sequence. EREFinder uses a sliding window analysis, where the binding affinity (*K*_*d*_) of every sequential set of fifteen base pairs is calculated. EREFinder takes the inverse of each *K*_*d*_ value, resulting in *K*_d_^-1^ values, because larger values of *K*_d_^-1^ indicate stronger binding. This approach also increases the influence of small *K*_d_ values on the local mean, increasing the likelihood of detecting canonical EREs. The mean of all inverse *K*_*d*_^-1^ values is then calculated across the user-specified width of the sliding window. In addition to designating the width, the user can also choose the slide interval for the sliding window. At one extreme, EREFinder can provide the ER-binding affinity for every overlapping 15-base pair subsequence present in a fasta file, if the user selects a width of fifteen and slide interval of one. The output from EREFinder is given as a comma-delimited text file suitable for additional analysis in a spreadsheet program or a statistical package, such as R.

## Methods

We wished to test the performance of EREFinder to that of two other freely available software packages, MATCH™ (Kel et al. 2003.) and Cluster-Buster (Frith et al. 2003). As a test dataset, we chose the nuclear genome of *Homo sapiens* GRCh38.p9 because of the completeness of the human genome and the availability of a list of known estrogen responsive genes in humans (Bourdeau et al. 2004). While we did use EREFinder to analyze the entire human genome, we compared EREFinder to other software packages by focusing on Chromosome 19. The program MATCH™ was utilized through the TRANSFAC^®^ public release online (Matys et al. 2003). We chose the default matrix for ERE as well as the default search parameters online that would limit the number of false positives. The website would only allow an upload of 100,000bp, so the chromosome was broken into 196 parts with each part uploaded separately; we then downloaded the separate results and joined into a single file. We also evaluated Cluster-Buster, because, like EREFinder, it is capable of identifying dense clusters of motifs. We used the default ERE matrix in Cluster-Buster along with the default search settings.

EREFinder allows the user to determine how to approach the output. We chose to look for windows of high estrogen binding affinity (i.e., high *K*_*d*_^-1^ values) compared to the background as a means to compare runs within and between programs. Using a custom R script (supplemental materials) we removed any windows with less than 80% of the base pairs scored and we determined the mean *K*_*d*_^-1^ for all remaining windows in the sample. To determine if a window had a higher *K*_*d*_^-1^ than the mean, we chose a cut off of both three and five standard deviations above the mean. Our custom R script analyzed the trimmed input file, using a cut-off value, window size, and slide length size, and produced a file reporting the highest *K*_*d*_^-1^ values within each detected peak of ER-binding potential. This approach allowed a visualization of regions with high ER binding as well as the size of each region.

Next, we wished to test how each program performed at detecting EREs on Chromosome 19. Bourdeau et al. (2004) found ten estrogen responsive genes on Chromosome 19. We hypothesized that those genes would be more likely to have EREs proximal to the translation start site when compared to randomly selected genes. We randomly choose ten genes on Chromosome 19 for comparison. Using a custom R script, we compared the ERE detections from the software programs under consideration to the locations of the ER-responsive and random genes. While many studies have focused on regions extremely close to the translation start site (i.e., within 10,000 base pairs), Lin et al. (2007) have suggested activation can take place large distances upstream from the transcription start site or downstream of the stop site. We developed a custom script to use the output from EREFinder, Cluster-Buster or MATCH™ to determine the location of each ERE detection relative to gene locations. Our script also reported whether the ERE site was upstream, downstream, or within the gene of interest. We ran this pairing script for two different distances from the estrogen-responsive genes for all programs and settings (Table 1). We ran the same settings for the ten randomly selected genes. Finally, we chose the best performing settings for EREFinder and ran those settings across the entire human genome. Using the same methods as described, we then paired our peaks with 183 estrogen responsive genes from Bourdeau et al. (2004) and 183 randomly selected genes (Table 2).

**Table 1:**
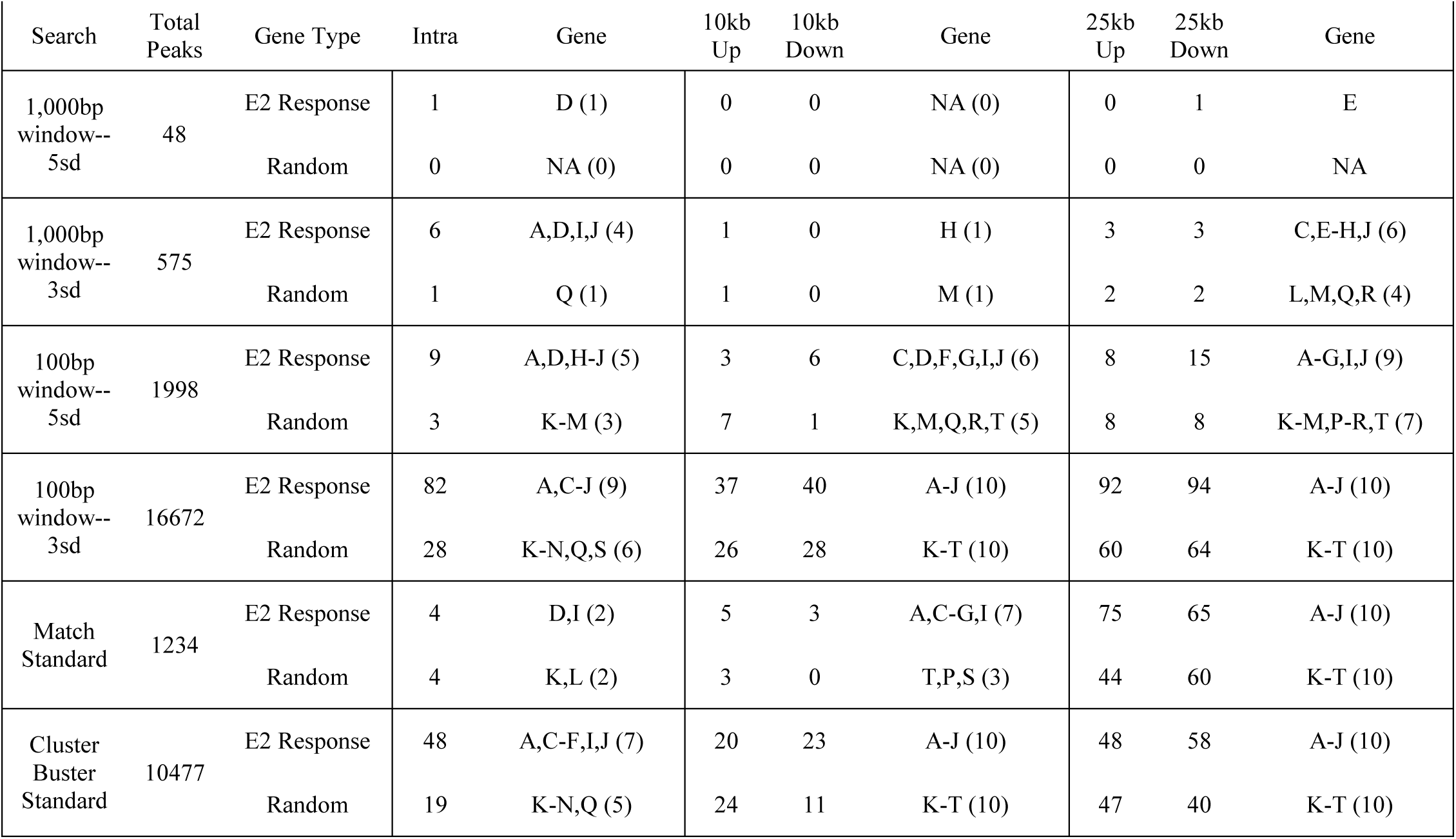
Number of Estrogen Receptor (ER) binding regions across human genome at chromosome 19 using EREFinder, MATCH™ (Kel et al. 2003.), and Cluster-Buster (Frith et al. 2003). EREFinder settings include size of window in base pairs and number of standard deviations (sd) above the mean used as the cutoff for high ER binding. Regions of high ER binding were matched with 10 known estrogen (E2) responsive genes and 10 random genes on the chromosome. Peaks were matched based on distance upstream from gene translation start site (Up), downstream from stop codon (Down), and in intragenic region (Intra). Genes found from the search are coded as follows with the number in parentheses indicating number of genes with a match. Estrogen Responsive Genes: A = C3, B = JUNB, C = PDE4C, D = ZNF91, E = USF2, F = SNRPA, G = EGLN2, H = ZNF230, I = CD33, J = ZNF17. Random Genes: K = CREB3L3, L = LOC400682, M = ZNF208, N = LINC00665, O = CYP2T3P, P = MIR4531, Q = LOC105372420, R = SNAR-F, S = ZSCAN5D, T = FKBP1AP1.

| Search | Total Peaks | Gene Type | Intra | Gene | 10kb Up | 10kb Down | Gene | 25kb Up | 25kb Down | Gene |
| --- | --- | --- | --- | --- | --- | --- | --- | --- | --- | --- |
| 1,000bp window--5sd | 48 | E2 Response | 1 | D (1) | 0 | 0 | NA (0) | 0 | 1 | E |
|  |  | Random | 0 | NA (0) | 0 | 0 | NA (0) | 0 | 0 | NA |
| 1,000bp window--3sd | 575 | E2 Response | 6 | A,D,I,J (4) | 1 | 0 | H (1) | 3 | 3 | C,E-H,J (6) |
|  |  | Random | 1 | Q (1) | 1 | 0 | M (1) | 2 | 2 | L,M,Q,R (4) |
| 100bp window--5sd | 1998 | E2 Response | 9 | A,D,H-J (5) | 3 | 6 | C,D,F,G,I,J (6) | 8 | 15 | A-G,I,J (9) |
|  |  | Random | 3 | K-M (3) | 7 | 1 | K,M,Q,R,T (5) | 8 | 8 | K-M,P-R,T (7) |
| 100bp window--3sd | 16672 | E2 Response | 82 | A,C-J (9) | 37 | 40 | A-J (10) | 92 | 94 | A-J (10) |
|  |  | Random | 28 | K-N,Q,S (6) | 26 | 28 | K-T (10) | 60 | 64 | K-T (10) |
| Match Standard | 1234 | E2 Response | 4 | D,I (2) | 5 | 3 | A,C-G,I (7) | 75 | 65 | A-J (10) |
|  |  | Random | 4 | K,L (2) | 3 | 0 | T,P,S (3) | 44 | 60 | K-T (10) |
| Cluster Buster Standard | 10477 | E2 Response | 48 | A,C-F,I,J (7) | 20 | 23 | A-J (10) | 48 | 58 | A-J (10) |
|  |  | Random | 19 | K-N,Q (5) | 24 | 11 | K-T (10) | 47 | 40 | K-T (10) |

**Table 2:**
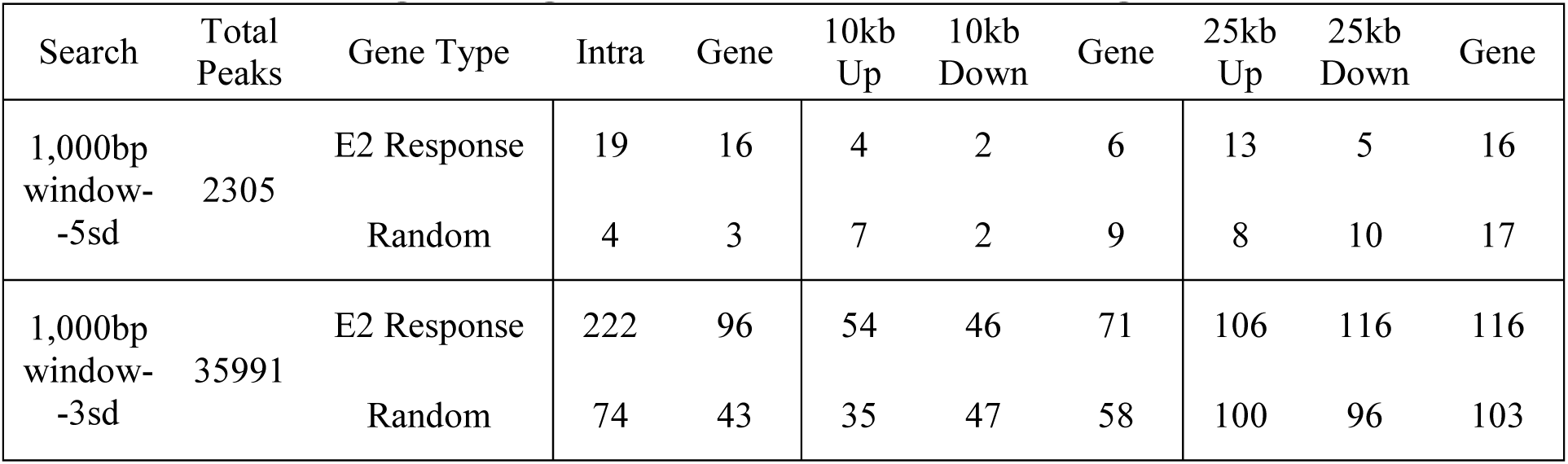
Number of Estrogen Receptor (ER) binding regions detected across the human genome using EREFinder. EREFinder settings include the size of the sliding window in base pairs and the number of standard deviations (sd) above the mean used to define high ER binding. Regions of high ER binding were matched with 183 known estrogen (E2) responsive genes and 183 random genes from the genome. Peaks were matched based on distance upstream from gene translation start site (Up), downstream from stop codon (Down), and in intragenic region (Intra). Total number of genes matched is included.

## Results

### EREFinder

No matter which settings were used, more peaks were associated with estrogen responsive genes than random genes; this pattern was especially true when looking in the intragenic regions (Tables 1 and 2). Larger windows led to fewer peaks and less frequent gene associations. Conversely, smaller windows led to ubiquitous associations no matter if the gene was estrogen responsive or randomly selected. Increasing allowable distance from the gene also led to more pairings with peaks across all settings. The two settings that appeared to perform the best were a 1,000-base-pair window with a three-standard-deviation above the mean cut-off and a 100-base-pair window with a five-standard-deviation above the mean cut-off. Anything with larger window sizes was unable to pair frequently to estrogen responsive genes, while anything with smaller window sizes was unable to distinguish from randomly selected genes. The three-standard-deviation, 1,000bp window did not detect ERE binding sites for as many genes as the five-standard-deviation, 100bp window. Nevertheless, the former settings did find twice as many putative ERE binding sites associated with estrogen responsive genes compared to random genes. We suggest that the 1,000bp window with a three-standard-deviation cutoff for significance provides the best balance of detection and false positives for a first pass through a genome. Using R, we generated a visual output for our chosen setting showing peak locations in relation to the estrogen response genes (Figure 1) as well as a close look at one estrogen responsive gene (Figure 2). Running our chosen setting across the whole human genome produced a similar result to what was seen on Chromosome 19 (Arnal et al. 2017).

**Figure 1:**
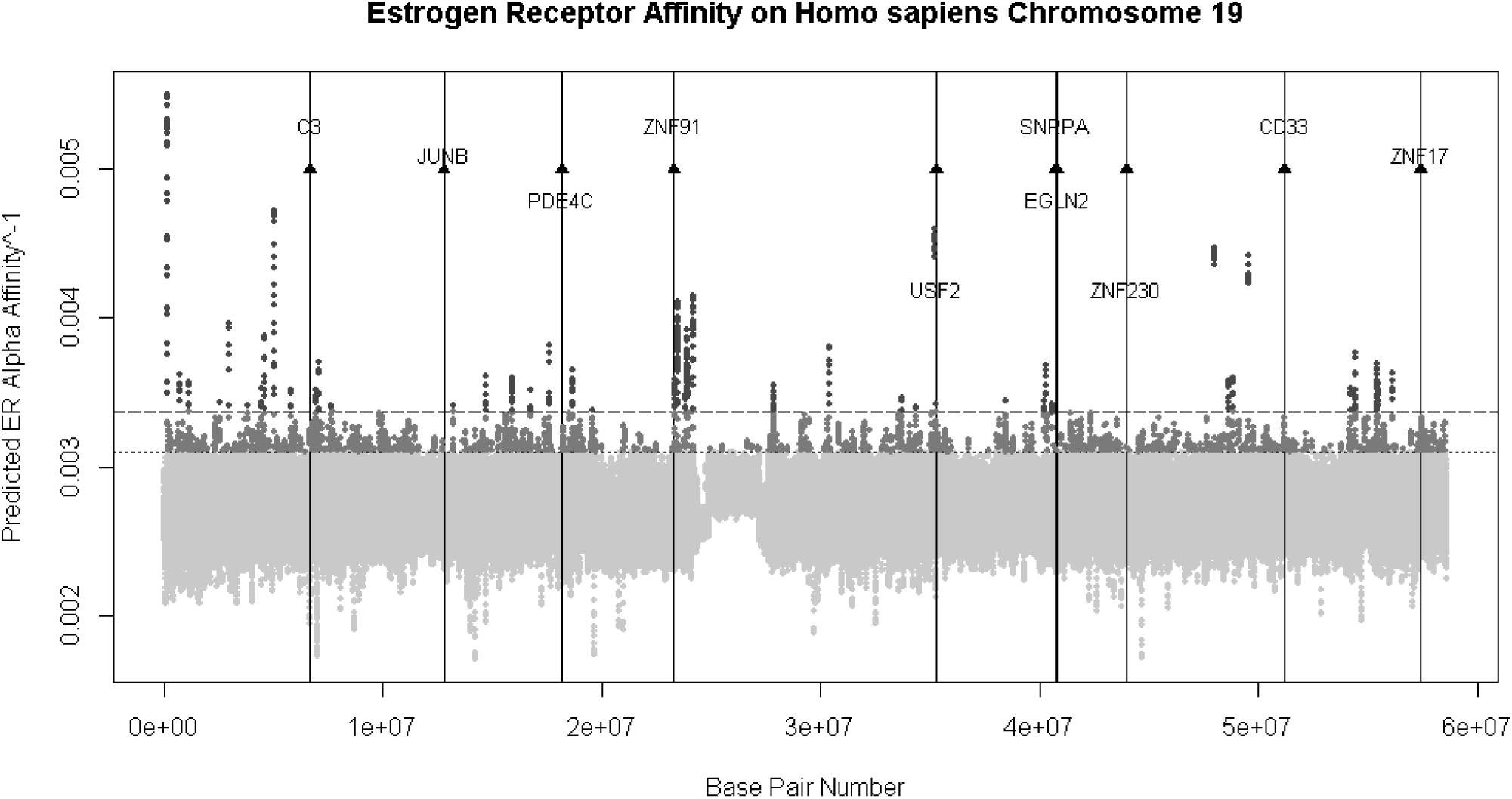
Output of EREFinder with a 1000 base pair window size sliding every 100 base pairs on Human Chromosome 19. Each dot represents a window’s ERα affinity, with grey dots representing values three standard deviations from the mean (99.7 percentile) and dark grey dots representing values five standard deviations from the mean (99.9999 percentile). Black vertical lines indicate locations of estrogen responsive genes on the chromosome.

**Figure 2:**
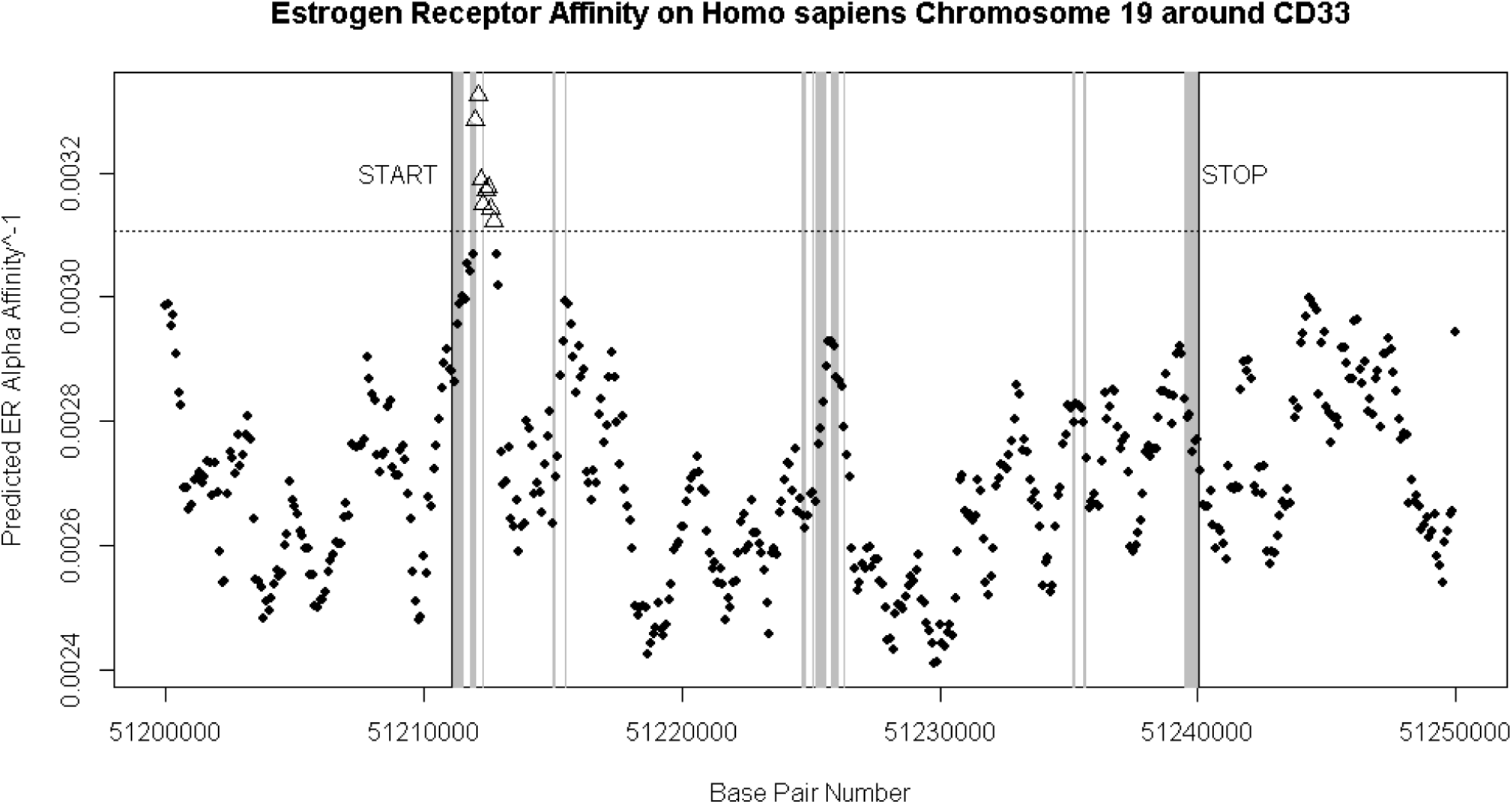
Output of EREFinder with a 1000 base pair window size sliding every 100 base pairs on Human Chromosome 19 around estrogen responsive gene CD33. Each dot represents a window’s ERα affinity with triangles representing values three standard deviations from the mean (99.7 percentile). Gray regions represent the exons.

### Comparisons to Other Programs

Cluster-Buster generated an excess of regions putatively responsive to estrogen receptors and had many regions proximal to the randomly selected genes, albeit in slightly lesser numbers than those proximal to estrogen responsive genes (Table 1). On the whole Cluster-Buster did not perform as well as EREFinder in this particular task. MATCH™ did better than Cluster-Buster and EREFinder when searches were limited to 10,000bp from the translation start site for distinguishing between estrogen responsive genes and randomly selected genes; however, EREFinder did better at distinguishing between the two in intragenic regions. At a 25,000bp limit, MATCH™ generated a result with all randomly selected genes paired to a hit. EREFinder, at the five-standard-deviation, 100bp window setting, performed comparably to MATCH™, except that EREFinder more strongly differentiated random genes from estrogen-responsive genes in the intragenic regions.

## Discussion

Here, we present a novel software package, EREFinder, which quickly scans whole genomes and calculates estrogen-receptor binding potential. Compared to other programs, EREFinder allows the user to see scores for every region considered and not just the ones determined through *a priori* settings. This feature provides more flexibility for user-defined searches and comparisons. While we have shown a technique for pairing regions of high ER binding with genes, this application is just one of many possible ways to manipulate the data for inquiry. Users could focus instead on the number of perfect half-sites within a window or locate perfect EREs in the sequence. EREFinder can be used both for *a priori* searches for ER binding in a given sequence (as demonstrated in this paper) and for *post hoc* searches around a gene of interest. These different types of searches facilitate the identification of regions of interest for hypothesis testing, comparative approaches, or evolutionary inquiries.

Our matching of EREs to estrogen-responsive genes found an increase in EREs in the proximal regions across all programs, indicating a general trend for EREs to be enriched on average near estrogen-responsive gene. Given the large number of hits and the occurrence of proximal EREs on random genes, we can be nearly certain that any bioinformatics approach to the detection of EREs will produce a large number of false positives. An ERE should be interpreted only as a putative location for ER binding, as studies have found that other factors that bind to DNA, such as FOXA1 (Carroll et al. 2005), are required to permit ER binding (Hah et al. 2013). Even if ER binding occurs, there must be a series of cofactors available to allow for transcription (Glass and Rosenfeld 2000, Shang et al. 2000). In addition, the interaction between ERα and ERβ can result in various expression patterns of up- and down-regulation, depending on the specific receptor that binds to the ERE (for review see Matthews and Gustafsson 2003).

With the variety of different factors that can affect expression and ER binding, it is important to consider that the primary function of EREFinder is to locate regions that have a sequence that could potentially bind an ER, without consideration of any other elements that may or may not be present on the given sequence.

While the presence of EREs is not necessarily an indication of ER binding, using EREFinder in conjunction with other empirical and computational methods could prove useful for a variety of research questions. EREs represent the physical location on the genome for ER binding, making EREs essential to the molecular process of ER gene activation and a useful point in the genome to search for putative ER binding. EREFinder demonstrates an ability to locate regions of high ER binding in a given sequence quickly and with an output that allows the user to manipulate the data in whatever manner they see fit.

